## Supplementary Materials for "Mapping epigenetic divergence in the massive radiation of Lake Malawi cichlid fishes"

### Supplementary Figures

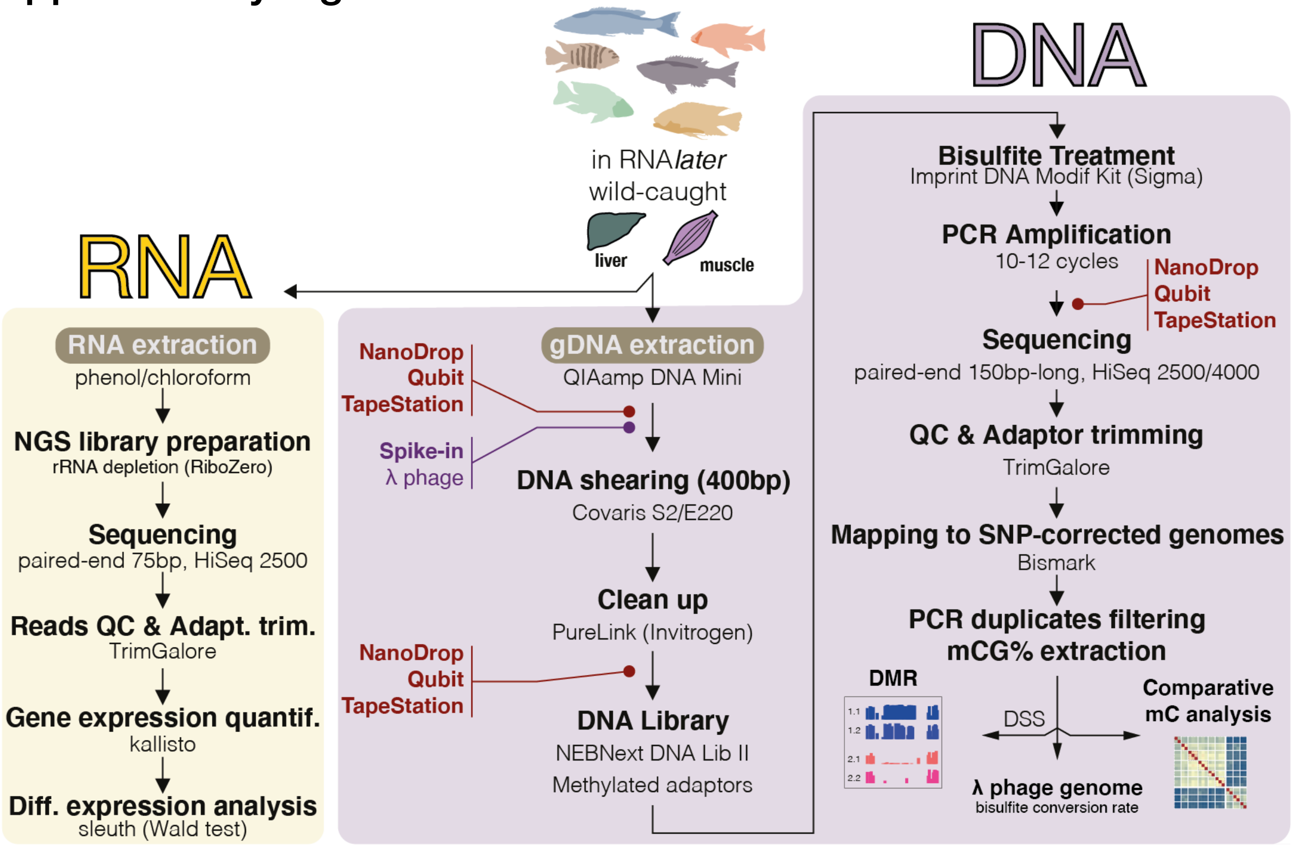

**Supplementary Figure 1. Experimental design.** Diagram of the main methods used to simultaneously generate total transcriptome sequencing (RNAseq) and whole-genome bisulfite sequencing (WGBS) datasets from liver and muscle tissues of the same wild-caught cichlid samples. Bulk whole-tissue sequencing was carried out to assess whole-tissue methylome and transcriptome variation between Lake Malawi cichlid species. See Fig.1c and Supplementary Table 1 for sample size information.

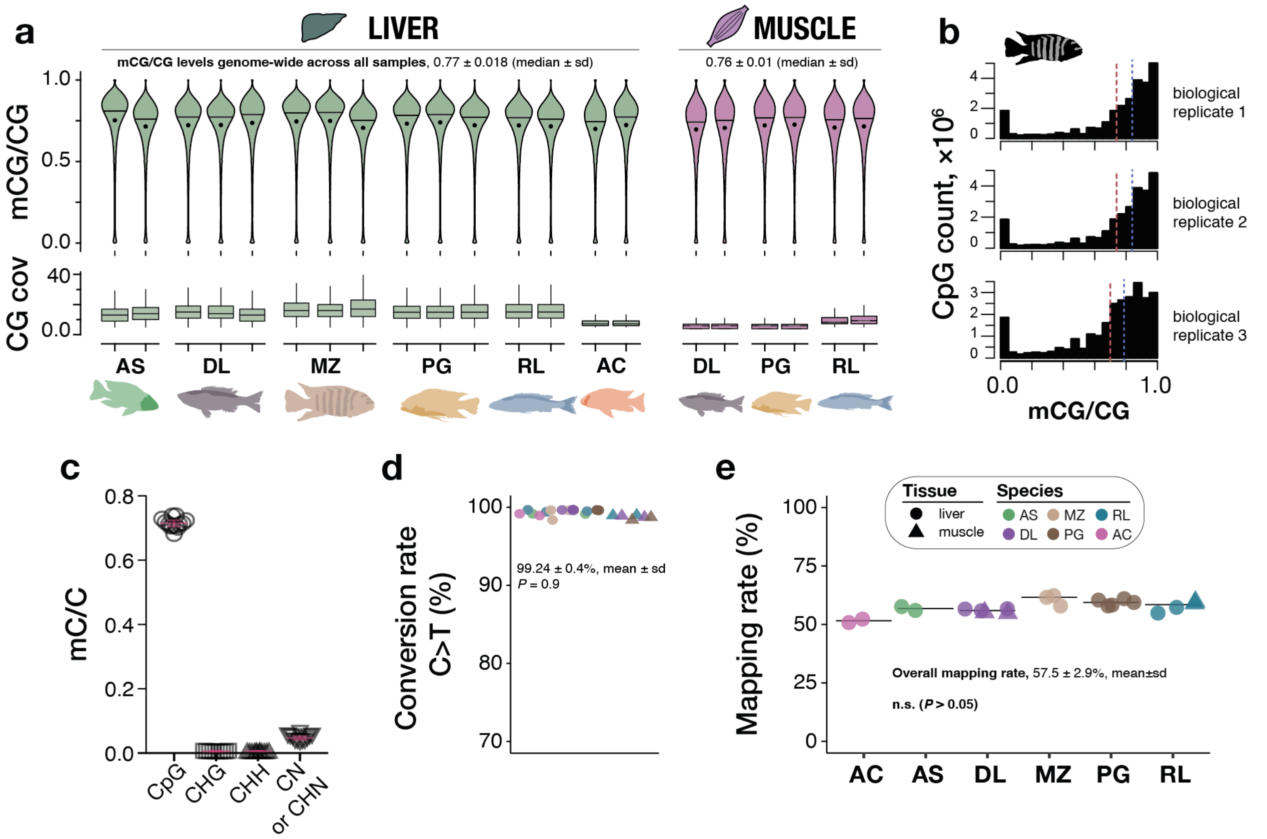

**Supplementary Figure 2. Genome-wide DNA cytosine methylation levels in Lake Malawi cichlids. a. Upper**: Violin plots showing the distribution of mCG/CG levels averaged in 1kbp windows genome-wide in both liver (green) and muscle (pink) tissues for all individual biological replicates per species. Black circles indicate mean values. Black lines indicate median values. **Lower**: boxplots showing sequencing coverage at all mapped CG sites for each biological replicate per species (only uniquely mapped, non-PCR duplicate reads shown, with ≥5 and ≤100 sequencing coverage at CG dinucleotides; 24.2 million CG dinucleotide sites). **b**. Bar plots of mC levels at individual mapped CG dinucleotides in the three *M. zebra* liver biological replicates. **c.** Percentage of methylated C (mC) over total C in different sequence contexts in all samples ($N$=21 data points). 25$^{th}$, 50$^{th}$ and 75$^{th}$ percentiles are indicated in red. H stands any base except for G; N for any base. **d.** Bisulfite conversion rates (%) were calculated in all samples using spiked-in unmethylated lambda DNA (See Methods and Supplementary Notes) and were not significantly different across samples (two-sided $P$ value for Kruskal Wallis test, $P = 0.9$, $N = 21$). **e**. Mapping efficiencies of all WGBS samples using species-specific SNP-corrected versions of the *Maylandia zebra* reference genome (UMD2a; see Methods; % of total paired-end 150bp-long reads). Black lines represent median values. Mapping rates were not significantly different across samples (two sided $P$ values for Dunn's Kruskal Wallis tests with Bonferroni multiple testing correction, two-sided $P > 0.09$). RL, *Rhamphochromis longiceps*; DL, *Diplotaxodon limnothrissa*; MZ, *Maylandia zebra*; PG, *Petrotilapia genalutea*; AS, *Aulonocara stuartgranti*; AC, *Astatotilapia calliptera* sp. Mbaka. WGBS samples: n = 3 biological replicates for liver DL, MZ and PG; n = 2 biological samples for liver AS, AC and RL, and muscle DL, PG and RL. All box plots indicate median (middle line), 25th, 75th percentile (box) and 5th and 95th percentile (whiskers) as well as outliers (single points)

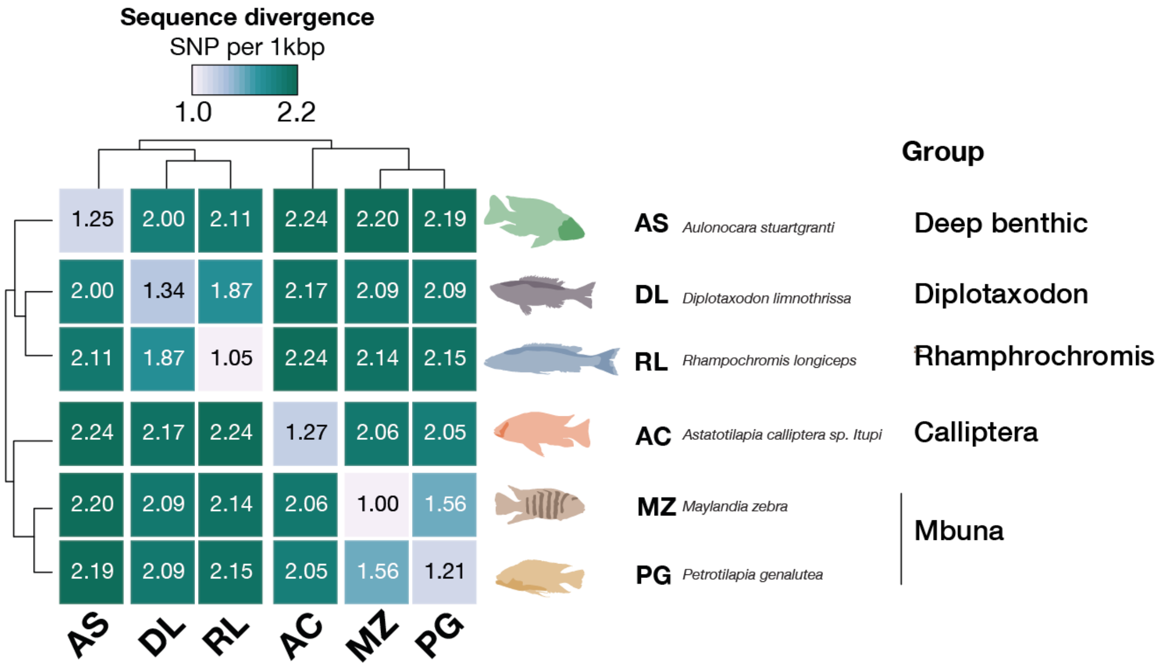

| | AS | DL | RL | AC | MZ | PG |
|---|---|---|---|---|---|---|
| AS | 1.25 | 2.00 | 2.11 | 2.24 | 2.20 | 2.19 |
| DL | 2.00 | 1.34 | 1.87 | 2.17 | 2.09 | 2.09 |
| RL | 2.11 | 1.87 | 1.05 | 2.24 | 2.14 | 2.15 |
| AC | 2.24 | 2.17 | 2.24 | 1.27 | 2.06 | 2.05 |
| MZ | 2.20 | 2.09 | 2.14 | 2.06 | 1.00 | 1.56 |
| PG | 2.19 | 2.09 | 2.15 | 2.05 | 1.56 | 1.21 |

**Supplementary Figure 3. Sequence divergence in Lake Malawi cichlid species part of this study.** Heatmap of pairwise genome-wide sequence divergences (SNP per 1kbp). Data from Malinsky et al. 2018.

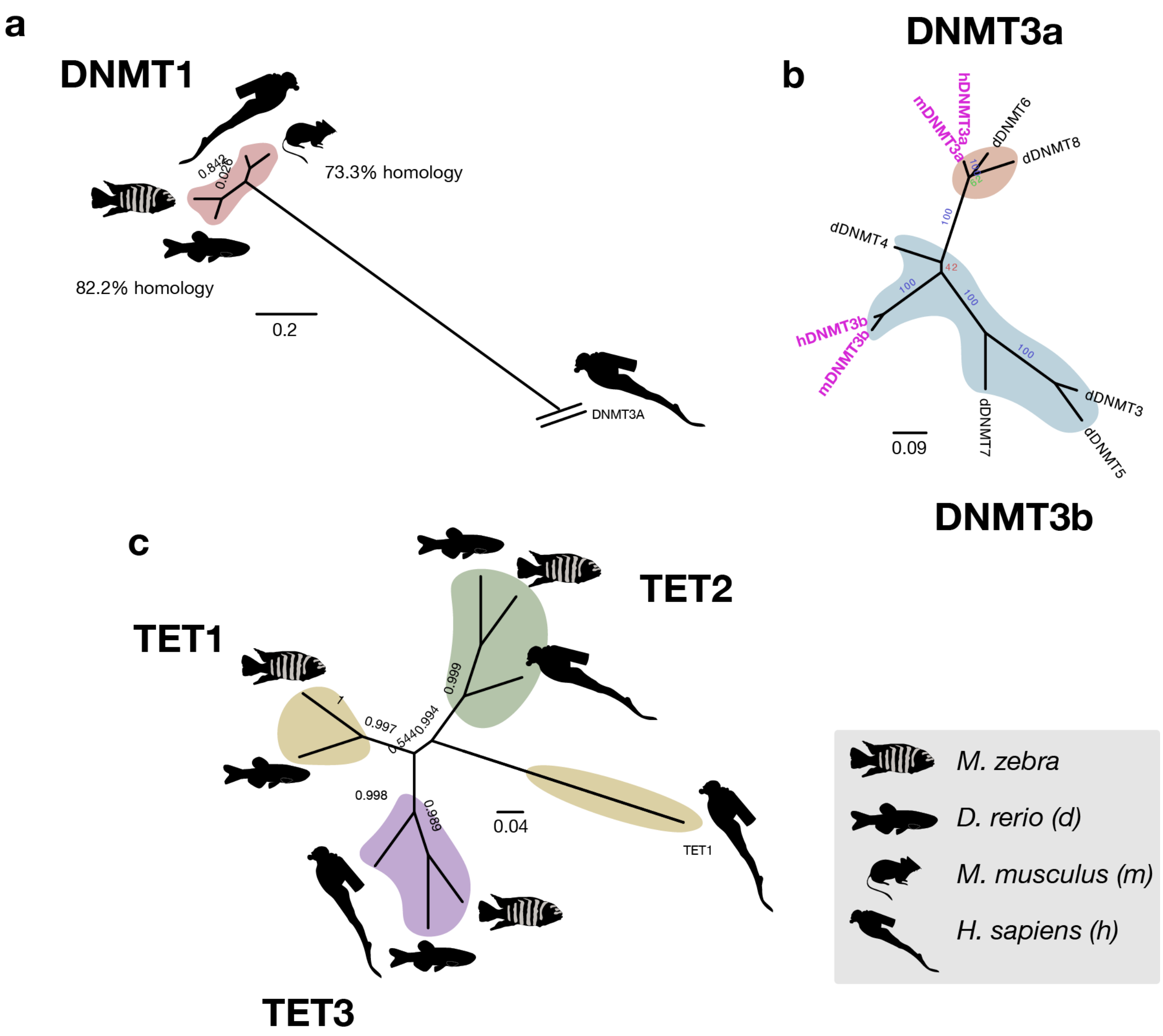

**Supplementary Figure 4.** Lake Malawi genomes has copies of principal DNMTs and TETs. Unrooted trees based on sequence homology with zebrafish, mouse and human for DNMT1 (**a.**), DNMT3a and DNMT3b (**b.**) and TET1-3 (**c.**).

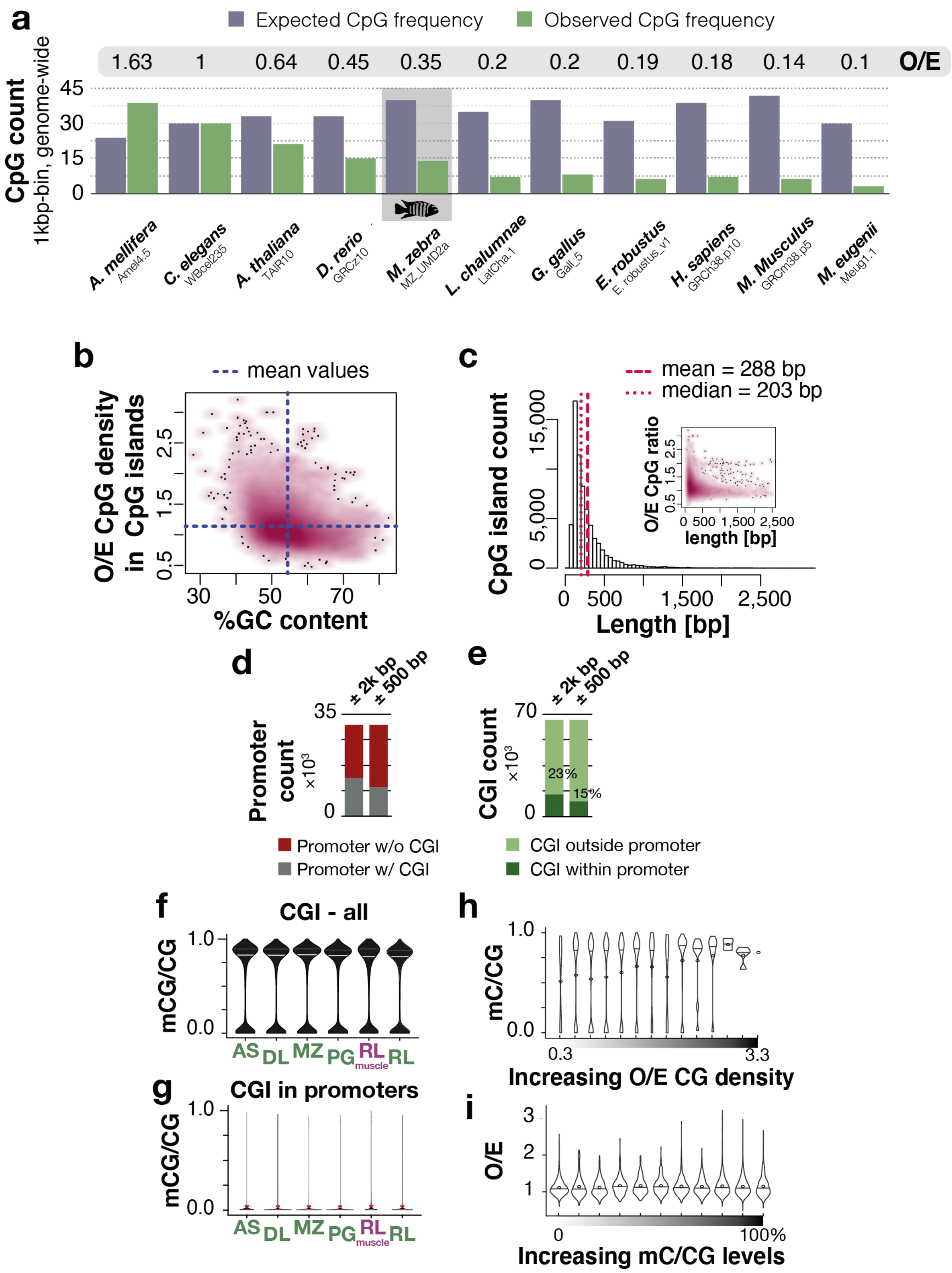

**Supplementary Figure 5. Predicted CpG islands (CGIs) in Lake Malawi genome. a**. Bar plots showing Observed/Expected (O/E) for CG content per kbp genome-wide across different species. O/E ratios shown above graphs. **b.** Scatter plot of CG content (%) against O/E at predicted CGIs (*n*=66,106) in MZ_UMD2a (MZ) reference genome. **c.** Histogram of CGI lengths in bp in MZ genome. Inlet graph, scatter plot showing CGI length (bp) against CpG density in CGI (O/E). Scatter plot colour density obtained through 2D kernel density estimate for b.-c. **d**. Count of promoter regions (TSS ± 500bp and TSS ± 2kbp) containing predicted CGIs in MZ genome. **e.** Count of predicted CGIs within promoters (either TSS ± 500bp or TSS ± 2kbp). **f.-g**. Violin plots of mC levels in all predicted CGIs (f.) and in CGIs within promoters (g.) in all samples (averaged by species; liver samples in green, muscle in purple) *n* = 3 biological replicates for livers of MZ, DL, PG; *n* = 2 biological replicates for AS and RL samples. **h.** mC levels in all CGIs according to CG frequency (increasing observed/expected CG content) in MZ liver (*n* = 3 biological replicates). **i.** Violin plots of CG content levels per mC level categories at predicted CGIs in MZ liver (*n* = 3 biological replicates). Horizontal lines and dots in all violin plots represent median and mean values, respectively

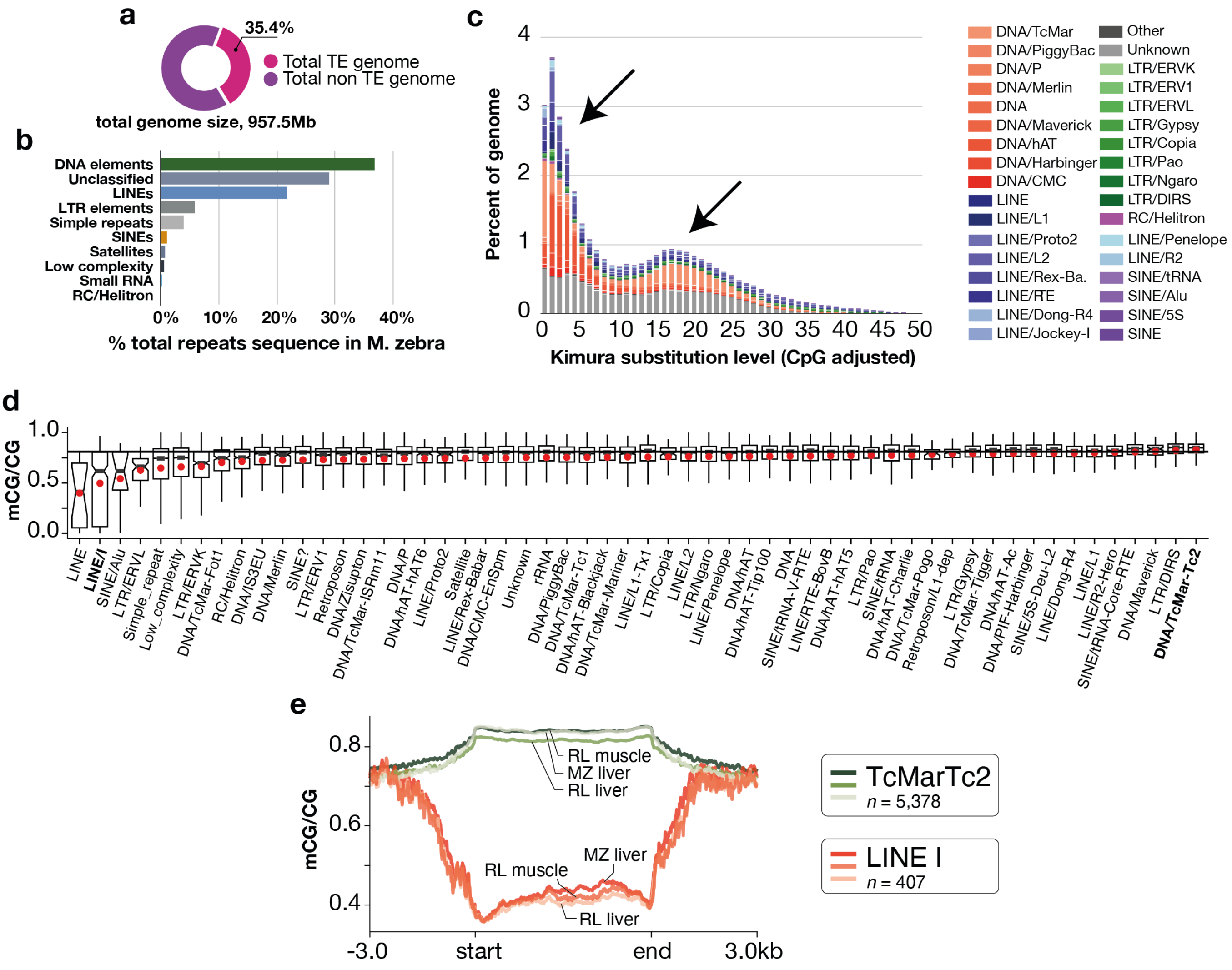

**Supplementary Figure 6. Genomic landscape of transposable elements and repeats in Lake Malawi cichlid *Maylandia zebra* UMD2a genome. a.** Proportion of TE (transposable elements and repeats) sequences predicted in *Maylandia zebra* (MZ) genome using RepeatModeler and RepeatMasker (see Methods). **b.** Percentage of total TE sequences for each TE category. **c.** Landscape of TEs in MZ genome. TEs are classified into known TE categories and according to sequence divergence from consensus sequences (Kimura substitution levels, analogous to TE age). Two distinct waves of burst of TE activity are indicated by black arrows. Graphs produced using RepeatModeler. **d.** Boxplot showing the average mC levels across transposable elements and repeat sequences predicted in MZ genome (average mCG/CG in $n$ = 3 biological replicates, liver). Red dots indicate mean values. Box plots indicate median (middle line), 25th, 75th percentile (box) and 5th and 95th percentile (whiskers) as well as outliers (single points). The black horizontal line indicates genome-wide average mC levels in MZ liver ($n$ = 3 biological replicates). **e.** Average mCG/CG levels over two examples of transposon subfamilies predicted in the genome of *M. zebra*, namely the DNA transposon Tc2-*mariner* (green) and the retrotransposon LINE I (orange) in muscle of RL (*Rhamphochromis longiceps*, $n$ = 2 biological replicates) and livers of both MZ ($n$ = 3 biological replicates) and RL ($n$ = 2 biological replicates). Counts ($n$) of genomic elements for each TE subfamily predicted in the genome of MZ indicated at the bottom of the legend caption.

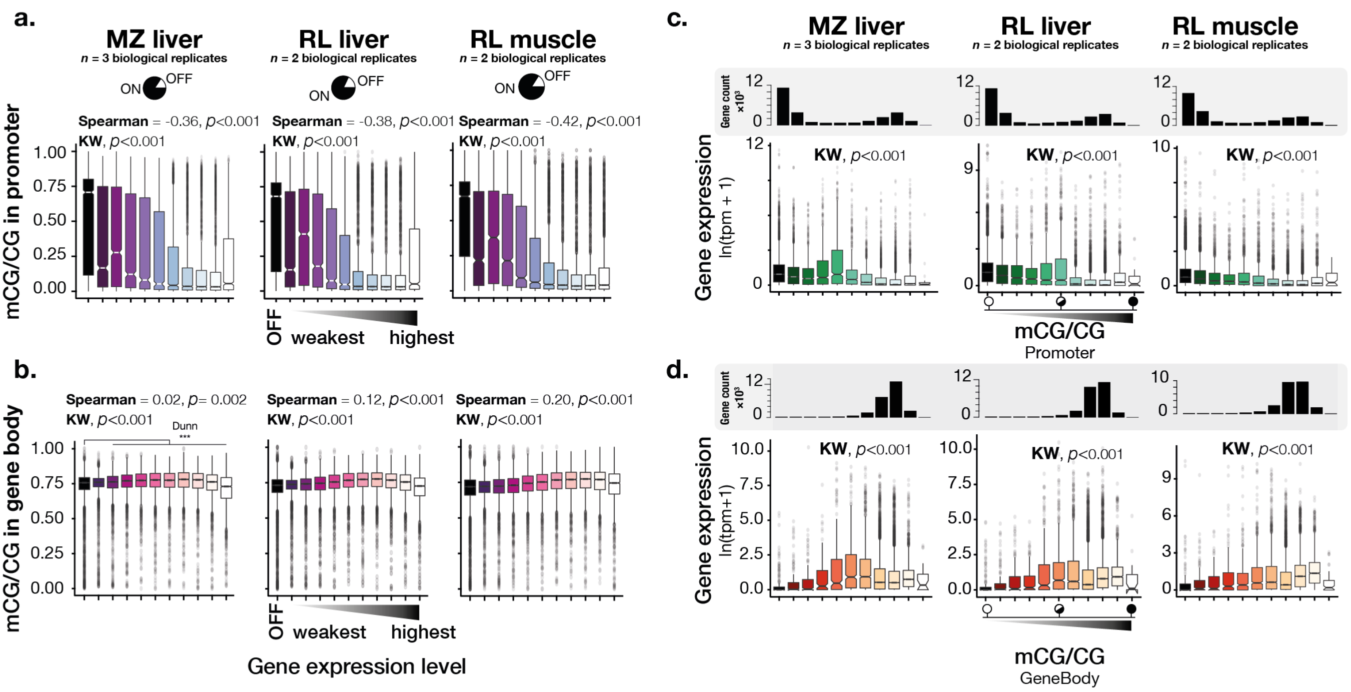

**Supplementary Figure 7. DNA methylation at promoter regions and gene bodies is significantly associated with differential gene expression activity. a, b.** Boxplots representing methylation levels (mCG/CG) at promoters (TSS ± 500bp; **a.**) and gene bodies (**b.**) according to gene expression levels in liver and muscle tissues. Genes were grouped into 11 categories (2,103 genes in all): one category for unexpressed/silent genes ("OFF") and 10 'ON' categories of increasing transcriptional activity (gene counts in each 'ON' category, *n*=2,103). Pie charts represent the number of silent/unexpressed ('OFF') and of expressed genes ('ON'). Gene expression levels are averaged transcript per million (TPM) values per gene per tissue per species. Methylation values are averaged mCG/CG ratios per loci per species (see below for *n* numbers). Scores for Spearman's rank correlation tests (between transcriptome activity and methylome level), as well as scores and two-sided *P*-values for Kruskal-Wallis (KW) followed by Dunn's multiple testing correction (between all categories) are indicated above boxplots (*P*<0.001 for all tests). **c, d.** Boxplots representing gene expression levels for different methylation level categories in promoters (**c.**) and gene bodies (**d.**). Two-sided *P*-values for Kruskal-Wallis followed by Dunn's multiple testing correction (KW; between all categories) are indicated above boxplots (*P*<0.001 for all tests). Promoters (**b.**) and gene bodies (**d.**) are grouped into categories based on their methylation levels (10 categories, from 0-100%; *n* values for number of genes in each category are shown in the bar plots above each graph series). x-axes represent averaged TPM values per species for each category. MZ and RL stand for *Maylandia zebra* and *Rhamphochromis longiceps*, respectively. For all graphs, *n* = 3 biological replicates for liver MZ (transcriptome and methylome datasets) and liver RL (transcriptome); n = 2 biological replicates for liver RL (methylome and transcriptome datasets). All box plots indicate median (middle line), 25th, 75th percentile (box) and 5th and 95th percentile (whiskers) as well as outliers (single points).

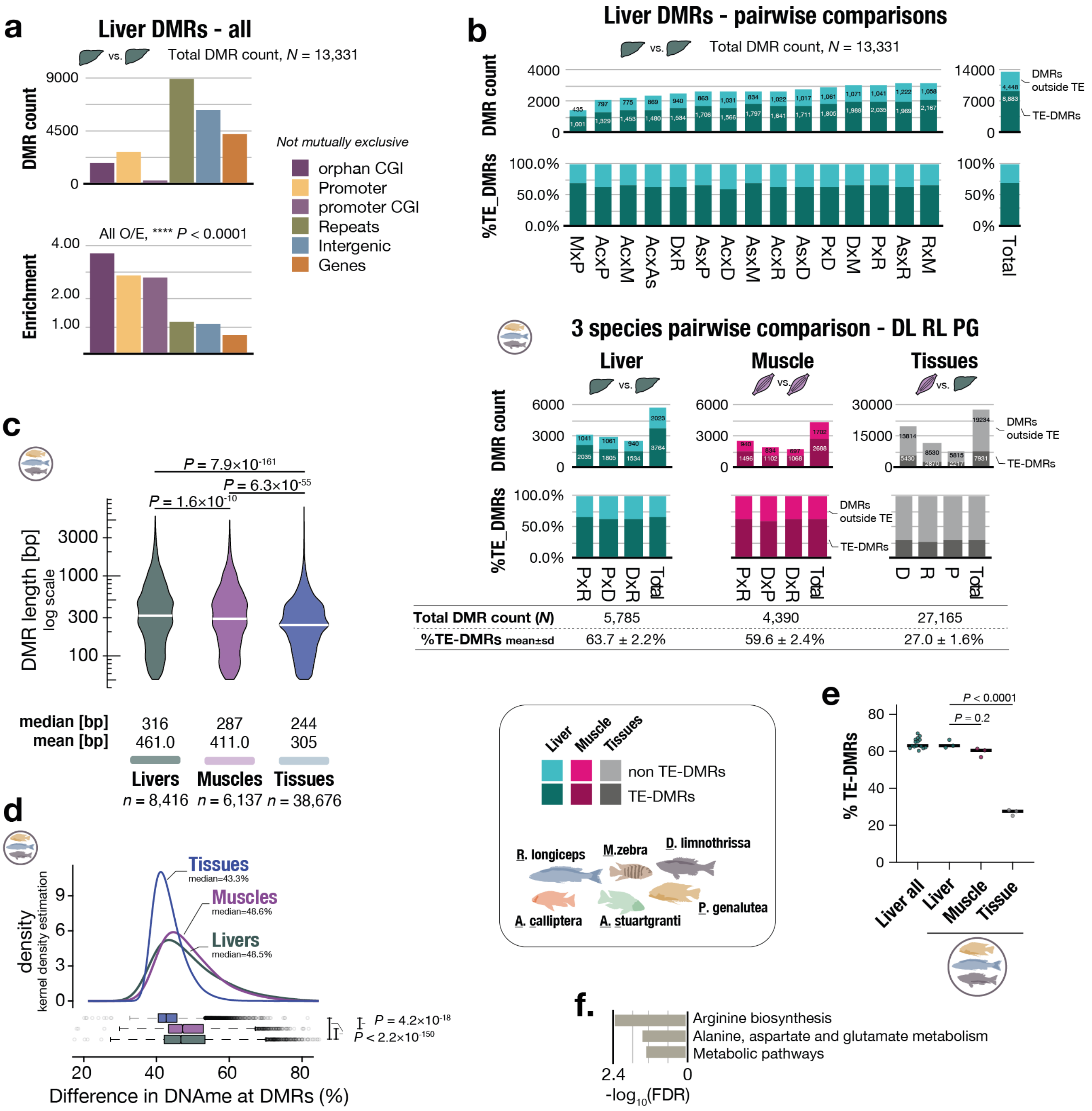

**Supplementary Figure 8. DMR count and enrichment of DMRs in different genomic locations. a.** Bar plots showing the total number of predicted between-species liver DMRs (upper graph) among the six cichlid species and the enrichment (observed/expected over chance; lower graph) in different genomic regions - non-mutually exclusive categories. DMRs with ≥25% overall methylation difference between >1 pairwise comparison ($p$<0.05) only are analysed further. CGI, CpG islands. $\chi^2$ tests, $P$ < 0.0001 for all O/E comparisons. **b.** Bar plots of the total number of DMRs predicted between each pairwise comparison for the six species (upper graphs). The proportion of DMRs for each species pairwise comparison localised in TE regions (TE-DMRs) is shown with a darker colour for each bar plot. Bar plots showing the percentage of TE-DMRs for each pairwise comparison (lower graphs). Lower panel shows the same plots for the three species comparison (RL, DL, PG) for which both muscle and liver WGBS data were available. **c.** Violin plots of the length (in bp) of all DMRs found between livers, between muscles and between tissues of the three species (RL, DL, PG). Median (white lines) and mean values of DMR length are given below each plot. Logarithmic scale for y-axis. Two-sided $P$-values for Dunn's Kruskal Wallis multiple tests (Bonferroni method) shown. **d.** Distribution plot (kernel density estimation) of mean DNA methylation difference (converted to absolute % values) between DMRs found between livers (n = 8,416), muscles (n = 6,137) and tissues (n = 38,876) of the three species (RL, DL, PG). Box plots indicate median (middle line), 25th, 75th percentile (box) and 5th and 95th percentile (whiskers) as well as outliers (single points), and are shown at the bottom (two-sided Bonferroni-corrected *P* values for Dunn's multiple tests shown for each comparison). Tissues vs Muscles, $P = 4.2\times10^{-18}$, Muscles vs Liver, and Tissues vs Livers, $P = 2.2\times10^{-150}$. **e.** Percentage of DMRs containing repeats/transposon elements (TE) for each DMR class (*P*-values for one-way ANOVA with Tukey's multiple comparison test shown above graphs). Black lines represent median values. **f.** Gene ontology enrichment analysis (KEGG terms only) for TE-DMRs located in promoter regions. $n = 3$ biological replicates for liver MZ, PG and DL; $n = 2$ biological replicates for liver AC, AS and RL, and muscle RL, DL and PG. Only GO terms with $P < 0.05$ (Benjamini-Hochberg false discovery rate [FDR]-corrected p-values) are shown.

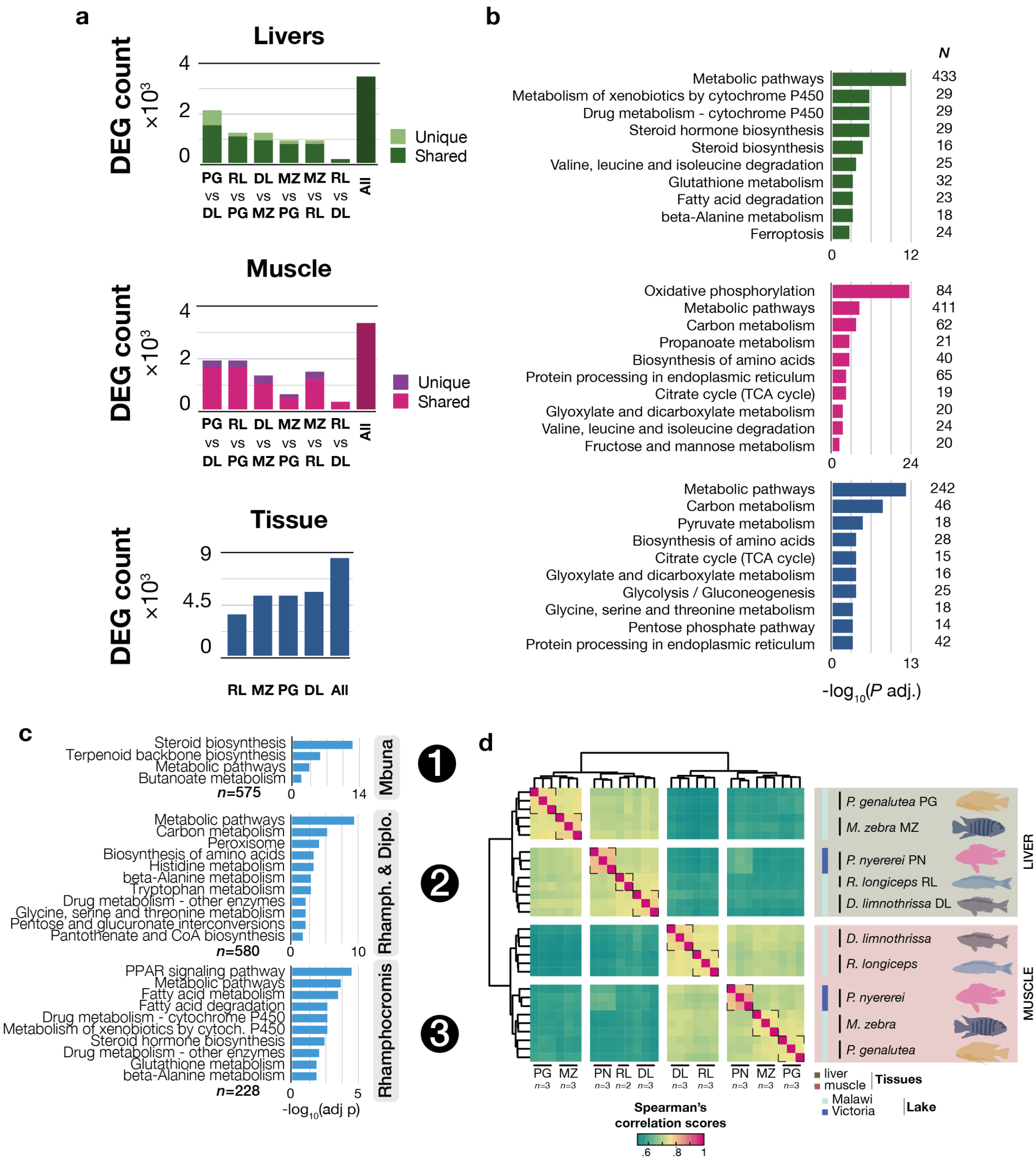

**Supplementary Figure 9. Transcriptome variation in Lake Malawi cichlids. a.** Total counts of differentially expressed genes (DEG) among liver (top; green), muscle (middle; red) and tissue (bottom; blue) transcriptomes for each pairwise comparison. Overall (All) counts shown for DEGs found in at least one pairwise comparison, $n$ = 3,437 for Livers; $n$ = 3,250 for Muscles; $n$ = 8,258 for Tissues; FDR < 0.01. **b.** GO enrichment analysis for liver (green, top), muscle (red, middle) and tissue (blue, bottom) DEGs - top 10 KEEG terms shown only. **c.** GO enrichment analysis for each DEG cluster shown in Fig. 3a. Only GO terms with $P$ < 0.05 (Benjamini-Hochberg false discovery rate [FDR]-corrected $P$-values) are shown. $N$ indicates the number of genes within each GO term category **d.** Heatmap and unsupervised hierarchical clustering based on whole transcriptome variation (using all annotated transcripts in *M. zebra* UMD2a) in both liver and muscle tissues of Lake Malawi cichlids. Spearman's rank correlation scores and Euclidean distances plotted (tree). Lake Victoria cichlid species, *Pundamilia nyererei* PN, was used as an outgroup species.

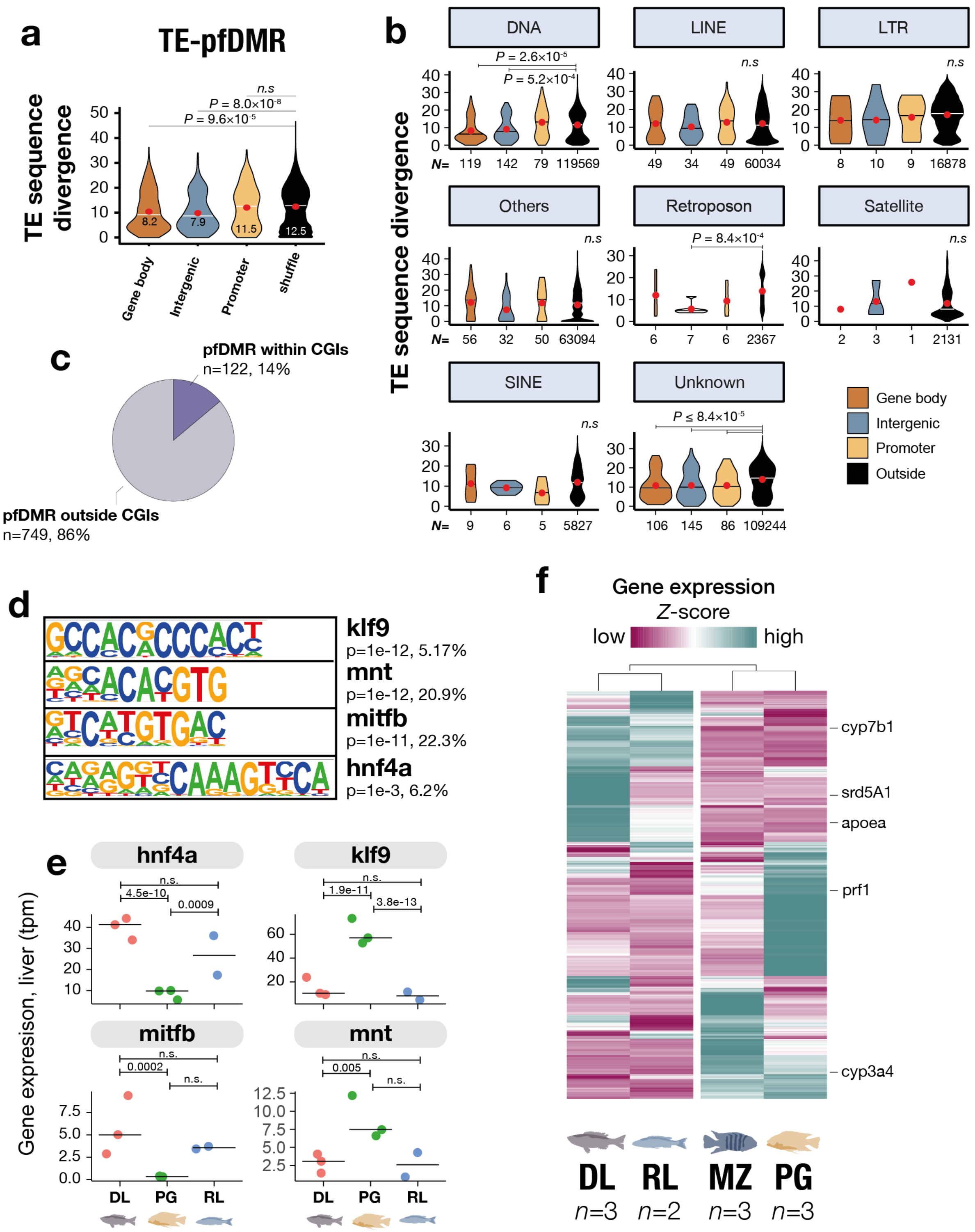

**Supplementary Figure 10. Putative functional (pf) DMRs are enriched in young transposable and repetitive elements (TE) and for liver-specific transcription factors binding sites, but not in CGIs. a**. Violin plots representing the distribution of TE sequence divergence (Kimura CpG-adjusted substitution levels; TE ages) for TEs associated with pfDMRs in each genomic feature (intergenic stands for intergenic regions, that is outside TSS ± 1kbp regions and gene bodies). White lines represent median values, red circles indicate mean values. Two-sided *P*-values for Kruskal–Wallis tests with Dunn's multiple comparison correction (Bonferroni method) are shown above each plot. Shuffle vs Gene body, $P = 9.6\times10^{-5}$, Shuffle vs Intergenic, $P = 8.0\times10^{-8}$. n.s., $P > 0.05$. **b.** Violin plots showing the distribution of TE sequence divergence (Kimura CpG-adjusted substitution levels; TE ages) for TE sequences associated with pfDMRs in each genomic feature and for each TE family. Two-sided *P*-values for Kruskal–Wallis tests with Dunn's multiple comparison correction (Bonferroni method) are shown above each plot. n.s., not significant ($P > 0.05$). pfDMRs are statistically associated with younger TEs in the case of DNA transposons (only for pfDMRs in intergenic regions and gene bodies, $P$ =

$5.2 \times 10^{-4}$ and $2.6 \times 10^{-5}$, respectively), retroposons (only for pfDMRs in intergenic regions, $P = 8.4 \times 10^{-4}$), and unknown/unclassified TEs (for pfDMRs in all the three genomic features, $P = 9.8 \times 10^{-8}$ [promoter], $P = 8.4 \times 10^{-5}$ [intergenic], $P = 3.2 \times 10^{-11}$ [gene body]).White lines represent median values, red circles indicate mean values. **c.** Pie charts summarising the proportion of pfDMRs localised in CpG islands (CGI). **d.** pfDMRs show significant enrichment for binding motifs (using HOMER) for differentially expressed TF genes in livers of Lake Malawi cichlid fish, with functions associated with hepatic gene expression regulation and liver functions, such as lipid homeostasis. klf9, krueppel-like factor 9; mnt, max-binding protein MNT; mitfb, microphthalmia-associated transcription factor; hnf4a, Hepatocyte nuclear factor 4 alpha. **e.** Graphs showing gene expression levels (transcript per million) in three Lake Malawi cichlid species for four TF whose binding motifs are enriched in pfDMRs. Two-sided *q* values (Bonferroni adjusted) for Wald tests shown above each graph (see Methods). Black lines represent median values. **g.** Heatmap of transcriptional activity in livers of genes associated with liver pfDMRs. z-score, averaged tpm values in liver for each species ($n = 767$). Examples of genes are indicated on the right. $n = 3$ biological replicates for liver MZ, PG and DL (RNAseq), $n = 2$ biological replicates for liver RL (RNAseq).

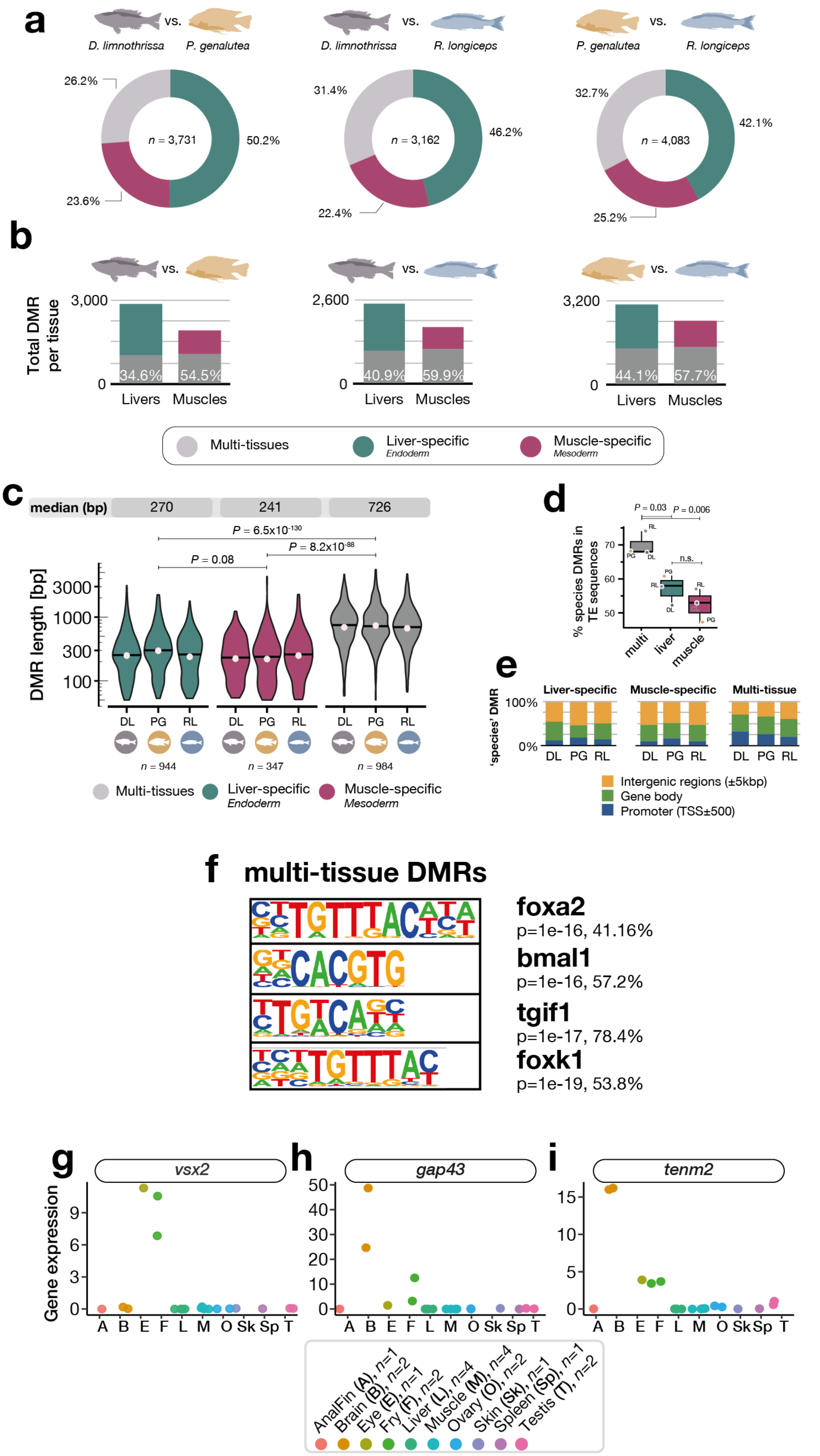

**Supplementary Figure 11. Tissue-specificity of between-species DMRs. a.** Pie charts summarising the type of DMRs found in each of the three pairwise species comparisons (% total). **b.** Bar plots summarising the absolute number of DMRs found between muscles (left) and livers (right) in each species pairwise comparison. Highlighted in grey are the DMRs found in both liver and muscle tissues (multi-tissue DMRs) within each tissue (percentage given within plot). DL in grey, PG in golden, RL in blue. DMRs with >25% overall methylation difference, $P < 0.05$. **c.** Violin plots showing the length in bp of DMRs associated with one respective species (compared to the others) for each category (liver only, muscle only and multi-tissues) found in *D. limnothrissa* (grey), *P. genalutea* (golden), *R. longiceps* (blue). x-axis, logarithmic scale. Median values of DMR length [bp] are given for each category (liver, muscle and multi DMRs) above the violin plots. Two-sided *P* values for Kruskal-Wallis tests corrected for multiple testing (Dunn's test, Bonferroni method) given for each comparison; $P = 0.29$ for muscle vs liver (n.s, $P = 0.08$). White circles and black lines represent mean and median values, respectively. DMR counts for each DMR category are given below the graph (*n*). Black lines represent median values, dots indicate mean values. **d**. Boxplots of the percentage of species-specific DMRs (over total species-specific DMRs) found in transposable elements (TE) for each DMR class. *P* values for ANOVA with Tukey's multiple comparison compared to multiDMRs are given above graphs (multi vs liver, $P = 0.03$; multi vs muscle, $P = 0.006$; muscle vs liver, $P = 0.6$ [n.s]). Box plots indicate median (middle line), 25th, 75th percentile (box) and 5th and 95th percentile (whiskers) as well as outliers (single points). **e.** Histograms of the proportion (in %) of species-specific DMRs for each genomic region (intergenic regions, gene body and promoters), for each DMR category (liver-, muscle-specific and multi-tissue). **f.** Multi-tissue DMRs found among the methylomes of the three cichlid species show significant enrichment for binding motifs (HOMER) for TFs with functions during liver development and embryogenesis. **g.-i.** Gene expression values for selected genes with multi-tissue DMRs (see Fig. 4e-g) in different tissues in *A. calliptera sp.* Itupi (n indicate biological replicates for each tissues). For WGBS dataset (average mCG/CG levels per tissue per species): $n = 2$ biological replicates for muscle RL, DL, PG and liver RL; $n = 3$ biological replicates for liver DL and PG.

### Supplementary Notes

##### Bisulfite conversion and WGBS read mapping

Bisulfite conversion was on average high (99.3 ± 0.4%, mean ± sd; Supplementary Fig. 2d and Supplementary Data 1). On average, using SNP-corrected genomes based on MZ_UMD2a reference genome assembly, 57.5 ± 2.9% (mean ± sd) of all sequenced reads were mapped uniquely (163.4 ± 29.8 million unique mapped paired-end reads, mean ± sd; see Supplementary Data 1).

##### CpG islands in Lake Malawi cichlid genome

Similar to other teleost fishes and, to a certain extent, mammals, the genome of *Maylandia zebra* is also highly CpG deprived overall (0.35 O/E CpG ratio; Supplementary Fig. 5a-b), yet features almost 66,000 CpG-rich regions (1.14 O/E CpG ratio; Supplementary Fig. 5c). These regions, typically 250bp in length (Supplementary Fig. 5c), display an overall strong binomial distribution of DNAme levels (Fig. 1d, Supplementary Fig. 5f). However, unlike mammals, only ~25% of cichlid promoters feature CGIs, which are almost exclusively unmethylated (Fig. 1d and Supplementary Fig. 5d, g). The majority of CGIs in cichlids lie outside promoter regions and are called 'orphan CGIs' (Fig.1 d and Supplementary Fig. 5d-f). The methylation level at all predicted CGIs is not correlated with CG content at CGIs (Supplementary Fig. 5h, i).

### Supplementary Tables

| | WGBS | | RNAseq | |
|---|---|---|---|---|
| | liver | muscle | liver | muscle |
| *Aulonocara stuartgranti* | 2* | 0 | 0 | 0 |
| *Maylandia zebra* | 3 | 0 | 3 | 3 |
| *Rhamphochromis longiceps* | 2 | 2 | 3 | 3 |
| *Petrotilapia genalutea* | 3 | 2 | 3 | 3 |
| *Diplotaxodon limnothrissa* | 3 | 2 | 3 | 3 |
| *Astatotilapia calliptera* sp. Mbaka | 2 | 0 | 0 | 0 |

**Supplementary Table 1.** Sample size for all Lake Malawi cichlid species part of this study. Only adult male specimens in full nuptial body colouration. * denotes that one specimen was female.

| | | *M. zebra* | *R. longiceps* |
|---|---|---|---|
| **Promoter** | **liver** | -0.36 | -0.38 |
| | **muscle** | n/a | -0.42 |
| **Gene body** | **liver** | 0.02 | 0.12 |
| | **muscle** | n/a | 0.2 |

**Supplementary Table 2.** Spearman's rank correlation scores (rho) between DNA methylation levels in promoter regions (TSS ± 500bp) and gene bodies (starting 500bp downstream TSS) and gene expression activity in both liver and muscle tissues for the two most phenotypically divergent Lake Malawi cichlid species of this study, namely *Maylandia zebra* and *Rhamphochromis longiceps*. $n$ = 3 for MZ liver, $n$ = 2 otherwise. $P$ values < 0.002 for all Spearman's rank correlation tests.
